# Specific Deletion of Axin1 Leads to Activation of β-Catenin/BMP Signaling Resulting in Fibular Hemimelia Phenotype in Mice

**DOI:** 10.1101/2022.07.04.498726

**Authors:** Rong Xie, Dan Yi, Qiang Jie, Qinglin Kang, Zeng Zhang, Zhenlin Zhang, Guozhi Xiao, Lin Chen, Liping Tong, Di Chen

## Abstract

Axin1 and Axin2 are key regulators of canonical Wnt signaling pathway. The roles of Axin1 and Axin2 in skeletal development and in disease development have not been fully defined. Here, we reported that Axin1 and Axin2 are essential for lower limb development. Specific deletion of *Axin1* in limb mesenchymal cells leads to fibular hemimelia (FH)-like phenotype, associated with tarsal coalition. Further studies demonstrated that FH disease is associated with additional defects resembling to the proximal femoral focal deficiency (PFFD) in *Axin1/Axin2* double knockout (KO) mice. We then provided *in vivo* evidence showing that Axin1 controls limb development through both canonical β-catenin and BMP signaling pathways. We demonstrated that inhibition of β-catenin or BMP signaling could significantly reverse the FH phenotype in mice. Together, our findings revealed that integration of Wnt and BMP signaling by Axin1 is required for lower limb development. Defects in Axin1 and Axin2 signaling could lead to the development of FH disease.

## Introduction

Fibular hemimelia (FH) is a congenital longitudinal limb deficiency characterized by complete or partial absence of the fibular bone. Unilateral fibular deficiency occurs in two-thirds of patients, with the right fibula being more often affected. FH may vary from partial absence of the fibula (10% of cases) with relatively normal-appearing limbs, to absence of the fibula with marked shortening of the femur, curved tibia, bowing of the leg, knee joint and ankle instability and significant soft tissue deficiency. The major functional deficiency results from limb length discrepancy in patients with unilateral FH or asymmetrical dwarfism in patients with bilateral FH. The foot is generally in an equinovalgus position. As there is limited growing potential within the affected bone, the extent of the deformity tends to increase with growth.

Occasionally, FH is associated with congenital shortening of the femur. Although it was first described by Gollier in 1698, the etiology of FH remains unknown (Stanitski et al., 2003). The deformity of FH is probably due to disruptions during the critical period of embryonic limb development, between 4th and 7th week of gestation. Vascular dysgenesis, viral infections, trauma and environmental influences have been suggested as possible causes. Most cases are sporadic. A family history has been reported in a small percentage of cases with an autosomal dominant pattern of inheritance and incomplete penetrance.

The evolutionarily conserved canonical Wnt signaling pathway controls many biological processes during the development and maintaining tissue homeostasis (Clevers et al., 2012). A key feature of this pathway is the regulation of its downstream effector β-catenin by a cytoplasmic destruction complex. Axin1 is a central scaffold protein of the destruction complex and directly interacts with all other core components in this complex (Clevers et al., 2012). It has been reported that Axin1 is the rate-limiting factor regulating β-catenin signaling (Lee et al., 2003). However, the *in vivo* role of Axin1 in the skeletal development and homeostasis has not been fully investigated due to early embryonic lethality (E9.5) of Axin1 mutant mice (Zeng et al., 1997). Genetic evidence from both humans and mice has implicated that Wnt/β-catenin signaling plays a crucial role in controlling all major aspects of skeletal development, including craniofacial, limb and joint formation (Baron et al., 2013; Regard et al., 2012). Bone morphogenetic protein (BMP) signaling also plays an important role in skeletogenesis during the development (Bandyopadhyay et al., 2006; Yoon et al., 2005). Thus, consistent with what is observed in many tissues and organs, Wnt and BMP signaling pathways have overlapped functions in controlling skeletal development and homeostasis. However, the key question is how the two pathways are integrated in controlling skeletal development and maintaining skeletal homeostasis.

Here we show that loss of *Axin1* in mouse limb mesenchymal cells resulted in severe defects in lower limb development, similar to FH disease phenotype and deletion of both *Axin1* and *Axin2* led to much severe FH phenotype associated with defects resembling to proximal femoral focal deficiency (PFFD). We found that inhibition of β-catenin signaling, either by deletion of one allele of *β-catenin* gene in limb mesenchymal cells or by the treatment with a specific *β-catenin* inhibitor, was able to significantly rescue the defects in FH phenotype observed in *Axin1* knockout (KO) mice. Furthermore, inhibition of BMP signaling also significantly reversed the defects in limb development and FH phenotype of *Axin1* mutant mice. Our findings indicate that Axin/β-catenin/BMP signaling plays a key role in FH development and pathogenesis.

## Results

### Deletion of *Axin1* in Prx1-expressing cells leads to FH-like phenotype

To determine the role of Axin1 in skeletal development and diseases, we generated *Axin1^Prx1^* KO mice by breeding the *Axin1^flox/flox^* mice (Xie et al., 2011) with *Prx1-Cre* transgenic mice (Logan et al., 2002) in which the Cre expression is under the control of the *Prx1* promoter. The *Prx1* (paired-related homeobox gene-1) regulatory element controls Cre expression throughout the early limb bud mesenchyme and in a subset of craniofacial mesenchyme (Logan et al., 2002). We examined skeletal development in E13.5 and E16.5 embryos and postnatal day 7 (P7) mice by Alizarin red/Alcian blue staining. The one notable defect is the presence of various fibular deficiencies in the *Axin1^Prx1^* KO embryos and postnatal mice (Figure 1A). The fibulae of *Axin1^Prx1^* KO mice did not mineralize even at day 7 postnatal stage (P7). Histological analysis using limb tissues dissected from *Axin1^Prx1^* KO mice showed that partially developed fibular tissues had high bone mass and poorly developed growth plate (Figure 1B). The number of chondrocytes was significantly reduced and the structure of growth plate was disorganized in *Axin1^Prx1^* KO mice (Figure 1B). To date, we have analyzed 52 *Axin1^Prx1^* KO mice and all of them have a fibular deficiency phenotype. Radiographic analysis of 4-week-old mice showed that the size of *Axin1^Prx1^* KO mice is smaller than Cre^-^ controls (Figure 1C). Radiographic analysis of 4- and 8-week-old mice also showed that some of *Axin1^Prx1^* KO mice were completely or almost completely absence of fibulae (>50% loss of fibulae, 27/52) where only a distal, vestigial fragment was present. The other *Axin1^Prx1^* KO mice had partial absence of the fibulae (30-50% loss of fibulae, 23/52) in which the proximal portions of the fibulae were absent while distal portions were present but could not support the ankle (Figure 1D, E). The mild fibular defect was observed in few of *Axin1^Prx1^* KO mice (2/52), in which the fibulae were absent less than 30% of their normal length (Figure 1F, G). These results demonstrate that Axin1 plays an essential role in fibular development. In addition to the fibular absence or hypoplasia, we also observed several additional defects in lower limbs of *Axin1^Prx1^* KO mice. All femora of *Axin1^Prx1^* KO mice were shorter and wider than those of their Cre^-^ littermates. The bowed tibia (Figure 1D-F) and the valgus ankle were observed and the tarsal coalitions (Figure 1H, I) were also found in 2-week-old *Axin1^Prx1^* KO mice. It is interesting to note that these skeletal defects observed in *Axin1^Prx1^* KO mice have been reported to be the key features of FH disease in humans (Achterman et al., 1979). In contrast, deletion of *Axin1* in *Sox9-*, *Col2-* and *Osx*-expressing cells does not affect fibular development (Figure 1J), suggesting that FH disease is caused by the defects in the specific cell population, limb mesenchymal cells.

**Figure 1.**
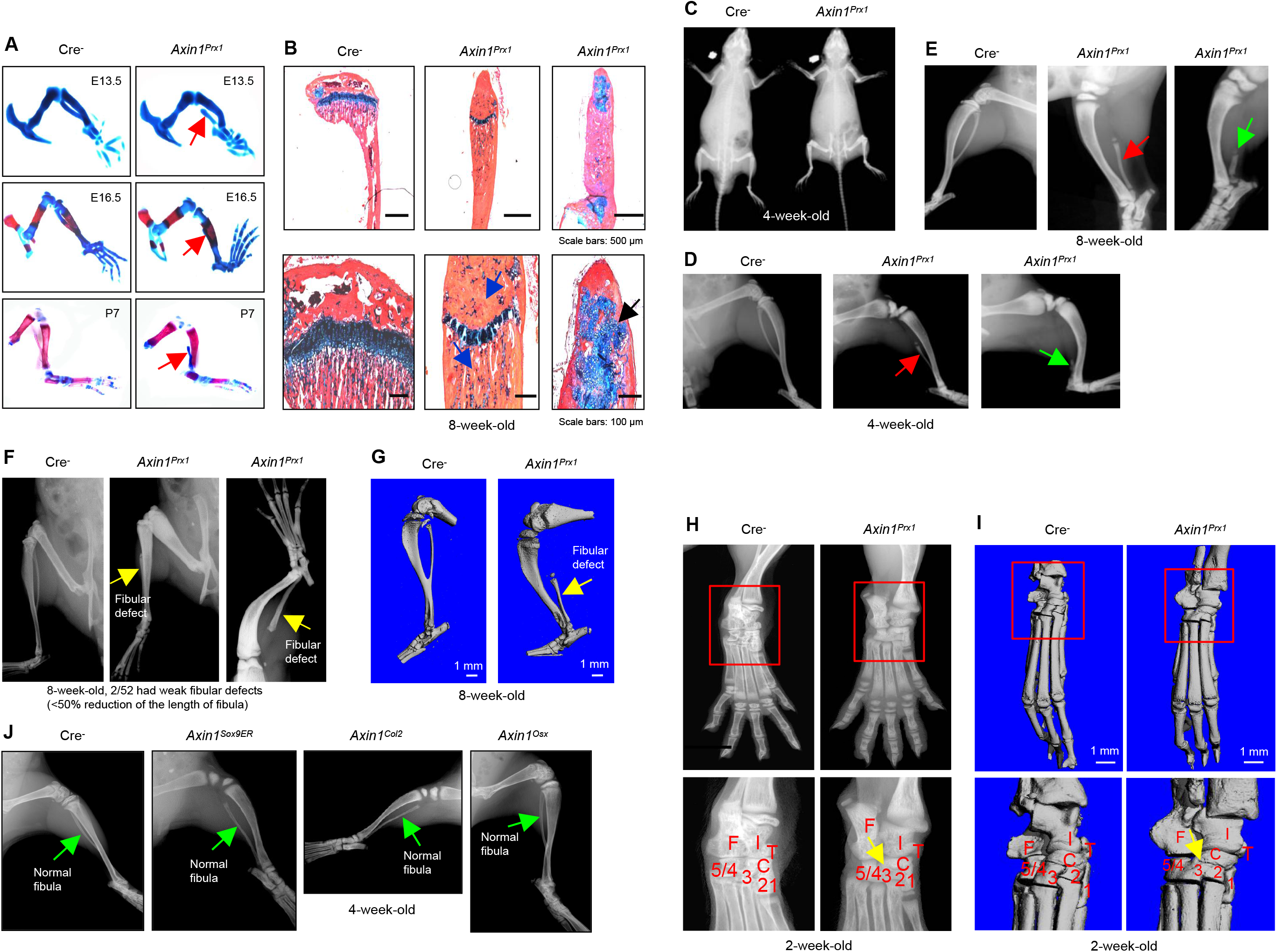
Deletion of *Axin1* in limb mesenchymal cells leads to defects resembling to FH disease. (**A**) *Axin1^Prx1^* KO embryos and postnatal mice showed partial development of fibula, which did not mineralize even at day 7 of postnatal stage (red arrows). (**B**) Histological analysis showed that disorganized fibular structure (black arrow), abnormal cartilage development and significant increased bone mass (blue arrow) were observed in 8-week-old *Axin1^Prx1^* KO mice. (**C**) Wholebody radiographic analysis revealed that size of 4-week-old *Axin1^Prx1^* KO mice is smaller than Cre^-^ control mice. (**D**) Defects in fibular development were observed in all 52 *Axin1^Prx1^* KO mice that we have analyzed. Radiographic analysis showed that the fibula in some of *Axin1^Prx^* mice was almost completely absent (>50% loss, 27/52) where only a distal, vestigial fragment was present (green arrow, right panel). The other *Axin1^Prx1^* mice had partial absence of the fibula (30-50% loss, 23/52) (red arrow, middle panel) in which the proximal portion of the fibula was absent while the distal portion was present but could not support the ankle. (**E**) Radiographic analysis also showed partial (red arrow) and almost complete absence of fibula (green arrow) in 8-week-old *Axin1^Prx1^* mice. **(F, G**) Radiographic and μCT analyses showed that fibulae were developed over 50% of their length in few *Axin1^Prx1^* KO mice (2/52). (**H, I**) Radiographic and μCT analyses also showed that fusion of tarsal elements, including 2nd, 3rd and 4th/5th tarsal elements and centrale, in hindlimbs was observed in 2-week-old *Axin1^Prx1^* KO mice (yellow arrow). (**J**) To determine the role of Axin1 in other cell populations, we generated *Axin1^Sox9ER^*, *Axin1^Col2^* and *Axin1^Osx^* conditional KO mice. X-ray radiographic analysis showed that deletion of *Axin1* in *Sox9-* and *Col2-* and *Osx*-expressing cells did not affect lower limb development.

### *Axin1^Prx1^*; *Axin2^+/-^* double KO mice have much severer defects in lower limb

Axin2 is the homolog of Axin1 and is 44% identical to Axin1 in their amino acid sequences. *Axin2* null mice (*Axin2^-/-^*) appeared craniofacial defects (Yu et al., 2005) and the high bone mass phenotype in adult *Axin2^-/-^* mice (Yan et al., 2009). Although *Axin2* mutant mice have no apparent fibular phenotype, deletion of *Axin1* in limb mesenchymal cells in combination with deletion of one allele of *Axin2* gene might result in more severe consequences for lower limb development. We then generated *Axin1^Prx1^*; *Axin2^+/-^* double KO (dKO) mice. The body size of the 3-week-old *Axin1/2* dKO mice is much smaller than *Axin1^Prx1^* KO mice (Figure 2A). And all *Axin1/2* dKO mice (n=15) displayed a complete absence of fibulae (Figure 2B). These results suggest that Axin2 also plays an essential role in fibular development. In addition, *Axin1/2* dKO mice had femoral defect phenotype. The femoral length was significantly shorter and the femoral head was not fully developed in 3-week-old *Axin1/2* dKO mice. No osseous connection between the femoral shaft and head and dysplasia of acetabulum were also found in *Axin1/2* dKO mice (Figure 2C). X-ray and μCT analyses showed complete missing of fibulae and hypoplasia of knee joint in 3-week-old *Axin1/2* dKO mice (Figure 2D). The tarsal coalition phenotype was much more severe in *Axin1/2* dKO mice and most tarsal bones, including calcaneus and distal tarsals 2, 3, 4, 5, were fused together (Figure 2E). In addition, *Axin1/2* dKO mice also exhibit knee joint fusion (Figure 2F). It is known that Wnt/β-catenin signaling regulates osteoclast formation. We performed TRAP staining and examined changes in TRAP-positive osteoclast numbers in *Axin1/2* dKO mice and found significant decreases in osteoclast formation in *Axin1/2* dKO mice (Figure 2G, H). Inhibition of osteoclast formation may contribute to the high bone mass phenotype observed in *Axin1/2* dKO mice. We also performed immunofluorescent (IF) staining of CD31 and VEGF and found that expression of CD31 as well as VEGF was significantly reduced in joint tissues of 3-week-old *Axin1/2* dKO mice (Figure 2I-L). Reduced CD31 and VEGF expression may be related to the hypoplasia phenotype of lower limb in *Axin1/2* dKO mice. Interestingly, the phenotype that we have observed in *Axin1/2* dKO mice resembles PFFD disease (Gillespie et al., 1983). We also analyzed changes in the gait of *Axin1^Prx1^* KO mice and *Axin1/2* dKO mice by taking videos. The video of WT mice showed normal walking gait (Video 1) and minor gait defect was observed in *Axin1^Prx1^* KO mice (Video 2). In contrast, severe gait abnormality was observed in *Axin1/2* dKO mice (Video 3). Hindlimbs were dragged by the forelimbs during walking in *Axin1/2* dKO mice (Video 3).

**Figure 2.**
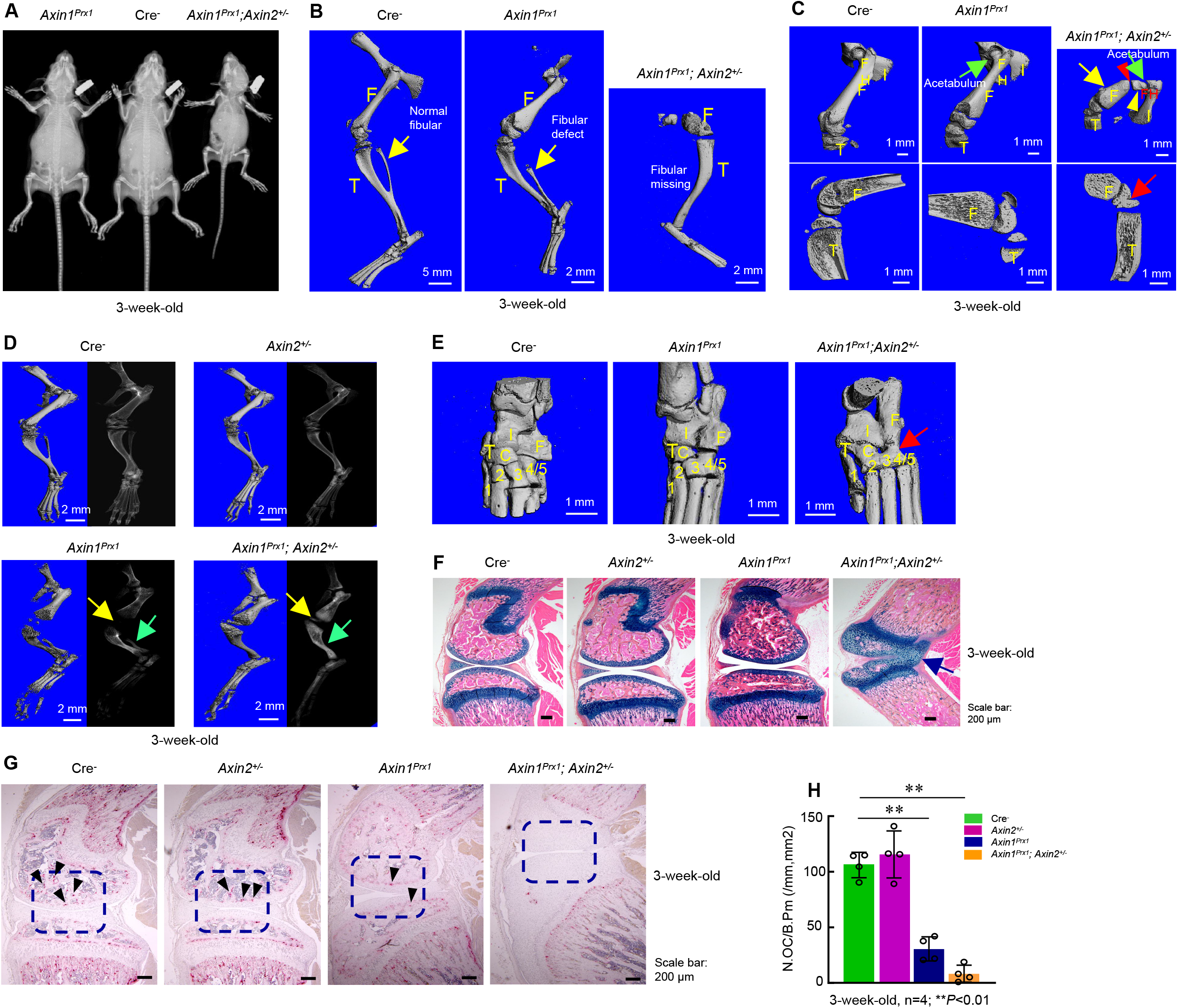
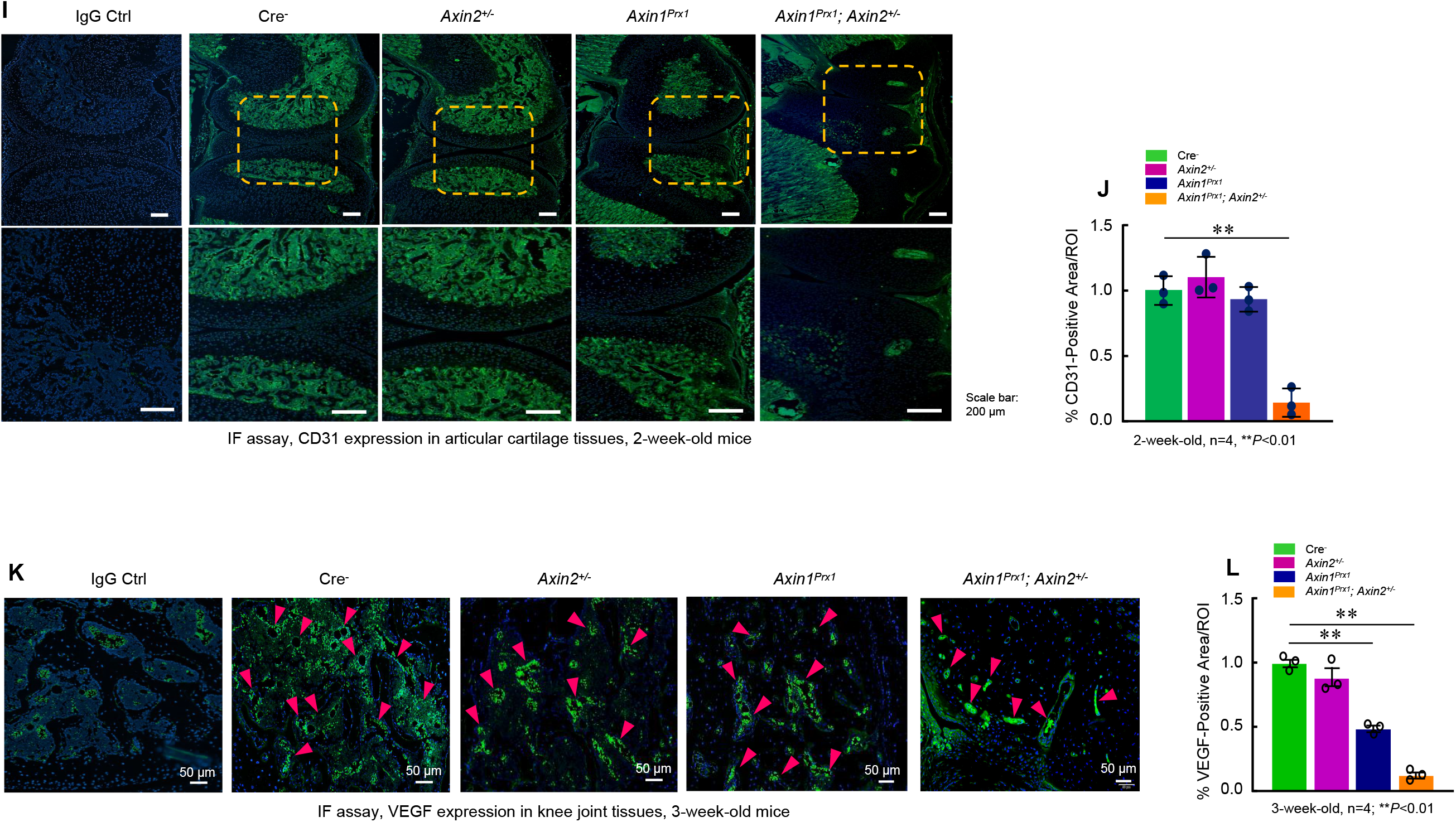
*Axin1^Prx1^*;*Axin2*^+/-^ double mutant mice (*Axin1/2* dKO mice) have much severe defects in lower limb development than *Axin1^Prx1^* KO mice. (**A**) X-ray radiographic analysis of 3-week-old mice showed that the size of *Axin1/2* dKO mice was much smaller than Cre^-^ control mice or *Axin1^Prx1^* KO mice. (**B**) We have analyzed 15 *Axin1/2* dKO mice and found fibulae were completely missing in all *Axin1/2* dKO mice. (**C**) μCT analysis showed that *Axin1*/*2* dKO mice (3-week-old) have features resembling to PFFD phenotype, including: (i) femoral shortening (yellow arrow); (ii) femoral head is not fully developed (yellow arrowhead); (iii) a large gap between the femoral head and proximal femoral shaft (red arrowhead); and (iv) defect in acetabulum formation (green arrow). F: femur; FH: femoral head; T: tibia; I: ischium. The knee joint fusion was observed in *Axin1/2* dKO mice (red arrow). (**D**) X-ray and μCT analyses showed that partial development of fibula in 3-week-old *Axin1^Prx1^* KO mice and completely missing fibula in 3-week-old *Axin1/2* dKO mice. (**E**) 3D images of μCT analysis of ankle joints showed that most of the tarsal elements of ankle joint were fused in *Axin1/2* dKO mice (red arrow, right panel). (**F**) Histology analysis also demonstrated the knee joint fusion in 3-week-old *Axin1/2* dKO mice. (**G, H**) We performed TRAP staining and found that reduction in osteoclast numbers in *Axin1^Prx1^* KO mice and much more severe defects in osteoclast formation in 3-week-old *Axin1/2* dKO mice. (**I-L**) Expression of CD31 and VEGF was analyzed in 3-week-old mice and significant decrease in VEGF expression was found in *Axin1^Prx1^* KO mice and severe reduction in both CD31 and VEGF expression was observed in *Axin1/2* dKO mice.

### Inhibition of β-catenin signaling reverses FH defects in *Axin1^Prx1^* KO mice

Since Axin1 is a well-known negative regulator of canonical Wnt pathway, deletion of *Axin1* will elevate β-catenin protein levels. If defects in fibular development in *Axin1^Prx1^* KO mice are due to elevated levels of β-catenin, reducing the expression levels of β-catenin may fully or partially correct defects observed in *Axin1^Prx1^* KO mice. To test this hypothesis, we examined genetic interaction between Axin1 and β-catenin during skeletal development in (*Axin1^flox/flox^*; *β-catenin^flox/wt^*)^*Prx1*^ double mutant mice. We found that deletion of one allele of the *β-catenin* gene under *Axin1^Prx1^* KO background significantly reversed defects in fibular development (Figure 3A, B) and caused reduction of BV from 92 to 71% (Figure 3C). In addition, we also used a specific β-catenin inhibitor iCRT14 to determine if blocking β-catenin signaling could reverse defects in the fibular development observed in *Axin1^Prx1^* KO mice. iCRT14 was shown to specifically inhibit β-catenin-induced transcription by disrupting the interaction between β-catenin and TCF4 (Gonsalves et al., 2011). The iCRT14 (2.5 mg/kg) was injected into *Axin1^Prx1^* KO mice (pregnant mothers at E9.5 stage, i.p. injection). The embryos were collected at E18.5. Histological analysis showed that the fibular defect phenotype in *Axin1^Prx1^* KO embryos were rescued by the treatment with iCRT14 (Figure 3D). The rescuing of fibular defects with iCRT14 treatment was also confirmed by μCT analysis in 4-week-old *Axin1^Prx1^* KO mice receiving iCRT14 treatment (Figure 3E). It is interesting to note that inhibition of Wnt secretion with LGK974 did not rescue the fibular defects observed in *Axin1^Prx1^* KO mice (Figure 3F). LGK974 is a specific small molecule compound of porcupine inhibitor, which inhibits Wnt secretion *in vitro* and *in vivo* (Liu et al., 2013). Together, these results demonstrate that Axin1 controls the fibular development through the canonical β-catenin signaling pathway.

**Figure 3.**
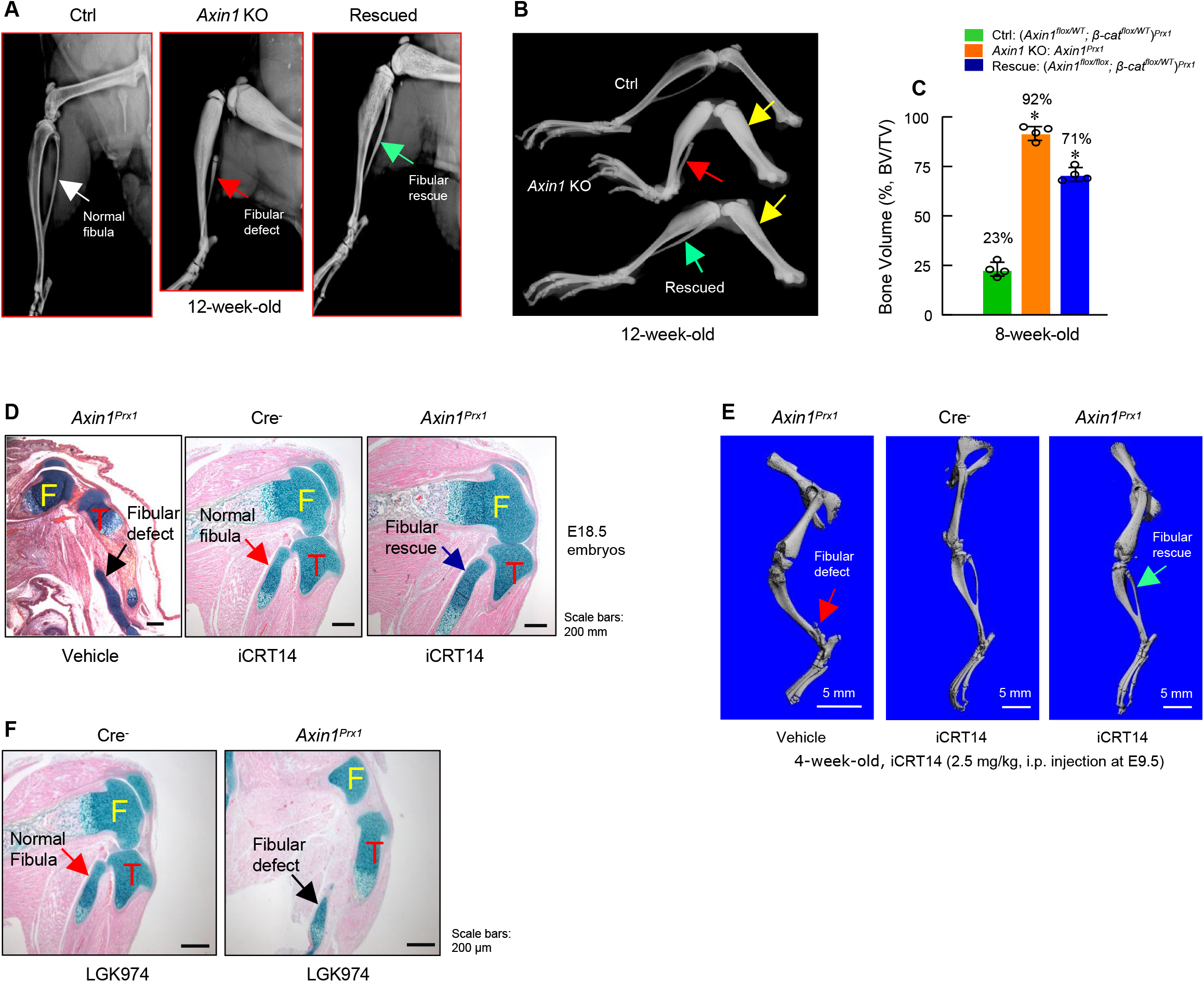
Inhibition of β-catenin signaling reverses defects in skeletal development observed in *Axin1^Prx1^* KO mice. (**A, B**) X-ray radiographic analysis showed that defects in fibular development observed in *Axin1^Prx1^* KO mice (red arrows) were significantly reversed (green arrows) by deletion of one allele of *β-catenin* gene (Rescued) (12-week-old mice). Yellow arrows showed bone mass was increased in *Axin1^Prx1^* mice and was partially reversed in Rescued mice. Radiographic analysis showed that deletion of one allele of *β-catenin* gene significantly reversed defects in fibular development in 12-week-old *Axin1^Prx1^* KO mice. (**C**) μCT analysis of hindlimbs of 8-week-old mice, including Ctrl: (*Axin1^flox/wt^/β-catenin^flox/wt^*)^*Prx1*^, *Axin1* KO: *Axin1^Prx1^*, and Rescued: (*Axin1^flox/flox^/β-catenin^flox/wt^*)^*Prx1*^ mice, showed that bone volume was reduced from 92% in *Axin1* KO mice to 71% in Rescued mice (n=4). (**D**) Results of histologic analysis of E18.5 embryos showed that treatment with β-catenin inhibitor, iCRT14 (2.5 mg/kg, i.p. injection to the pregnant mothers at E9.5 stage), almost completely reversed defects in fibular development in *Axin1^Prx1^* KO embryos. (**E**) μCT analysis confirmed that the treatment with iCRT14 reversed defects in lower limb development in *Axin1^Prx1^* KO mice, such as lack of fibula and bowed tibia. (**F**) In contrast, the treatment with LGK974 (inhibitor of Wnt secretion) failed to reverse FH phenotype observed in *Axin1^Prx1^* KO mice. Data presented in (c) were analyzed by one-way ANOVA followed by the Tukey Post-Hoc test (n=4, \**P*<0.05).

### Inhibition of BMP signaling reverses fibular defects in *Axin1^Prx1^* KO mice

In previous studies, we found that *Bmp2* and *Bmp4* expression was up-regulated in *Axin2* KO mice (Yan et al., 2009). To determine if BMP signaling is upregulated in *Axin1^Prx1^* KO mice, we extracted total RNA from hindlimbs derived from E12.5 Cre^-^ and *Axin1^Prx1^* KO embryos. We found that expression of *Bmp2*, *Bmp4*, *Gremlin1* and *Msx2* was significantly upregulated in limb tissues of *Axin1^Prx1^* KO embryos (Figure 4A-F). To determine if inhibition of BMP signaling will reverse fibular defects observed in *Axin1^Prx1^* KO mice, we injected *Axin1^Prx1^* KO mice with BMP signaling inhibitor dorsomorphin (2.5 mg/kg, i.p. injection) to the pregnant female mice at E9.5 stage. Dorsomorphin has been shown to inhibit BMPR-IA (ALK3), BMPR-IB (ALK6) and ALK2 activity (Yu et al., 2008). Analysis of histological sections of hindlimbs of E18.5 embryos showed that fibular defects in *Axin1^Prx1^* KO embryos were significantly rescued by the treatment with dorsomorphin (Figure 4G). The result of μCT analysis of 6-week-old mice confirmed that the fibular defect phenotype observed in *Axin1^Prx1^* KO mice was significantly reversed by the treatment with dorsomorphin (Figure 4H). The single dose of BMP inhibitor (2.5 mg/kg at E9.5 stage) is not able to reverse high bone mass phenotype caused by continuing upregulation of BMP signaling in *Axin1^Prx1^* KO mice. Also, dorsomorphin did not affect fibular development in the Cre^-^ control mice (Figure 4H). We also determined the stage specific effect of dorsomorphin on reversing fibular defects in 3-week-old *Axin1^Prx1^* KO mice and found that dorsomorphin lost its protective effect if injected after E12.5 stage (Figure 4I). Administration of dorsomorphin at E12.5 stage significantly reversed fibular defects as well as rescuing knee join dysplasia phenotype in 3-week-old *Axin1^Prx1^* KO mice (Figure 4J). In contrast, same concentration of TGF-β inhibitor, small chemical compound SB-505124, had no effect on reversing fibular defect in *Axin1^Prx1^* KO mice (Figure 4K). These results demonstrate that BMP signaling upregulation also contributes to the FH defects observed in *Axin1^Prx1^* KO mice.

**Figure 4.**
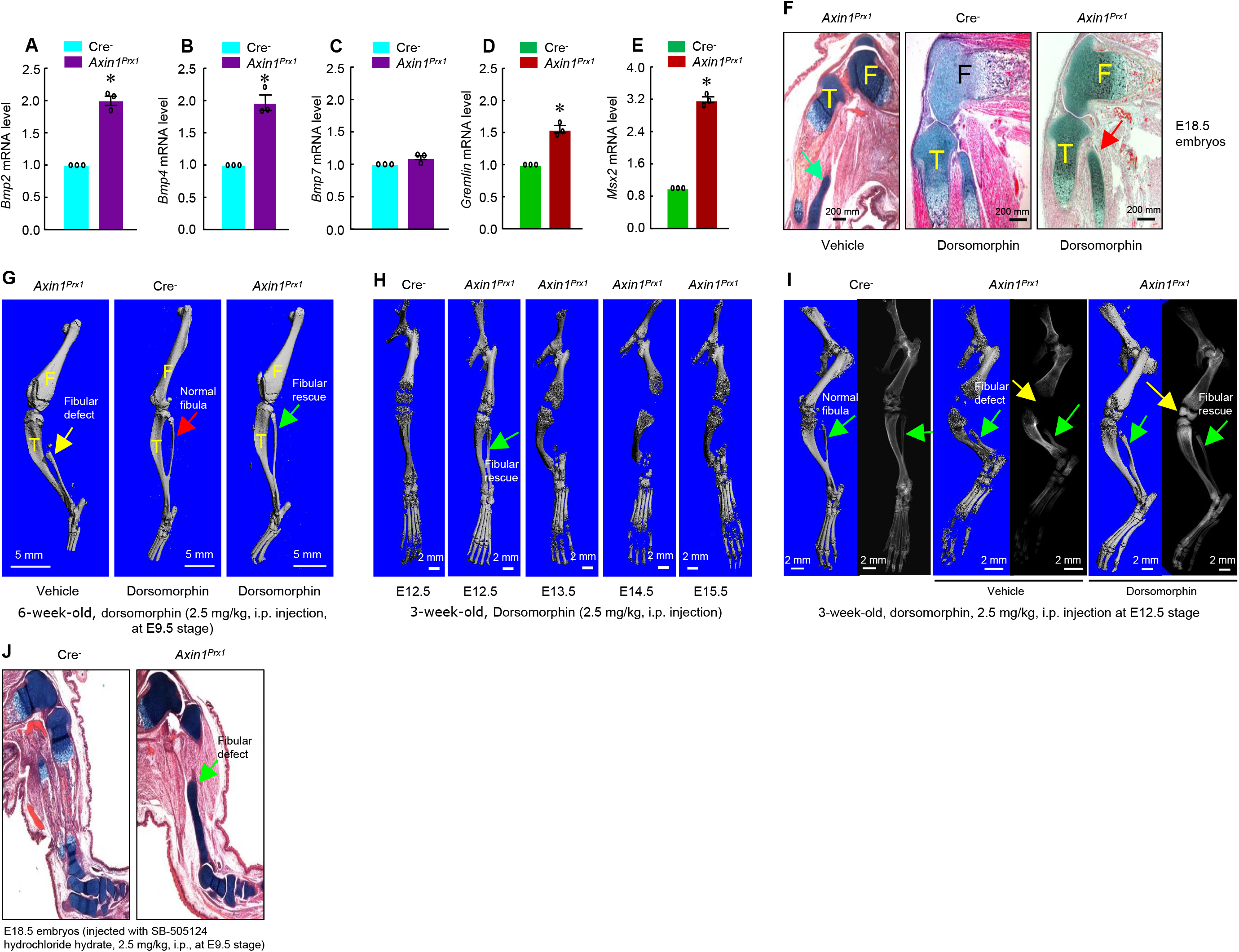
Inhibition of BMP signaling significantly reversed skeletal phenotype observed in *Axin1^Prx1^* KO mice. **(A-E**) BMP signaling was up-regulated in limb bud cells derived from E12.5 *Axin1^Prx1^* KO embryos. Real-time PCR analysis showed that expression of *Bmp2*, *Bmp4*, *Gremlin*, and *Msx2* was significantly up-regulated in limb bud cells derived from E12.5 *Axin1^Prx1^* KO embryos (\**p*<0.05, unpaired Student’s *t*-test, n=3). (**F**) Inhibition of BMP signaling by the treatment with BMP inhibitor, dorsomorphin (2.5 mg/kg, i.p. injection to E9.5 mothers), reversed fibular defects of *Axin1^Prx1^* KO embryos (E18.5). Green arrow (left panel): the defect in the fibular development. (**G**) μCT analysis of 6-week-old mice showed that the treatment with dorsomorphin (2.5 mg/kg, i.p. injection to E9.5 mothers) completely reversed defects in fibular development. (**H**) We also examined stage-specific effect of dorsomorphin treatment and found that dorsomorphin could effectively reverse defects in fibular development up to E12.5. No rescuing effect could be observed when dorsomorphin was administered at later stages, such as E13.5, E14.5 and E15.5. (**I**) Dorsomorphin could significantly reverse defects in fibular development when injected at E12.5 stage. (**J**) In contrast, injection of same amount of TGF-β inhibitor SB-505124 had no significant effect on fibular development.

### Axin1 inhibits BMP signaling through promoting pSmad5 degradation

Next, we sought to explore if Axin1 directly regulates BMP signaling although it is known that BMP signaling is downstream of β-catenin signaling in bone cells. Since Axin1 serves as scaffold protein, we examined whether there are interactions between Axin1 and Smad proteins. The results of co-immunoprecipitation (co-IP) assays revealed that endogenous Axin1 indeed interacted with Smad5 in C3H10T1/2 cells (Figure 5A). Then, we investigated whether Axin1 regulates the stability of pSmad5. In pulse-chase experiments, the limb cells from E12.5 Cre^-^ or *Axin1^Prx1^* KO embryos were treated with BMP2 for 0.5 hour, followed by incubation without BMP to track the levels of phosphorylated Smad5 (pSmad5) (Figure 5B, C). The pSmad5 levels decreased gradually after removal of BMP2 in Cre^-^ cells, but duration of pSmad5 was much longer in *Axin1* mutant cells (Figure 5B, C). The results indicate that Axin1 could also inhibit BMP signaling through promoting pSmad5 degradation. We next determined if the increased duration of pSmad5 by *Axin1* deletion is independent of β-catenin. We performed BMP2-induced pulse-chase experiments in the presence of iCRT14. The treatment with iCRT14 did not affect the duration of pSmad5 in Cre^-^ cells as well as in *Axin1*-deficient cells (Figure 5D). In contrast, LGK974 did block the Wnt3a-induced prolonged duration of pSmad5 (Figure 5D). These results suggest that Axin1 regulated pSmad5 levels are independent from β-catenin. To determine whether Wnt3a-induced pSmad5 increase occurs through inhibition of its degradation, serum-starved C3H10T1/2 cells were stimulated with BMP2 for 0.5 hour. After BMP2 was removed, the cells were treated with or without Wnt3a for 3 hours. Indeed, Wnt3a significantly prolonged the duration of the pSmad5 at the similar extend as the treatment with proteasome inhibitor MG132 (Figure 5E), and iCRT14 did not block the Wnt3a-induced prolonged duration of pSmad5 in C3H10T1/2 cells (Figure 5E). It is worth noting that total Smad5 levels did not change under these experimental conditions, suggesting that only a small fraction of Smad5 is phosphorylated in response to the treatment of BMP2 at the physiological level. These results are in agreement with previous observations (Fuentealba et al., 2007; Alarcon et al., 2009). Taken together, these data suggest that Axin1 could regulate BMP signaling through a direct and indirect mechanisms.

**Figure 5.**
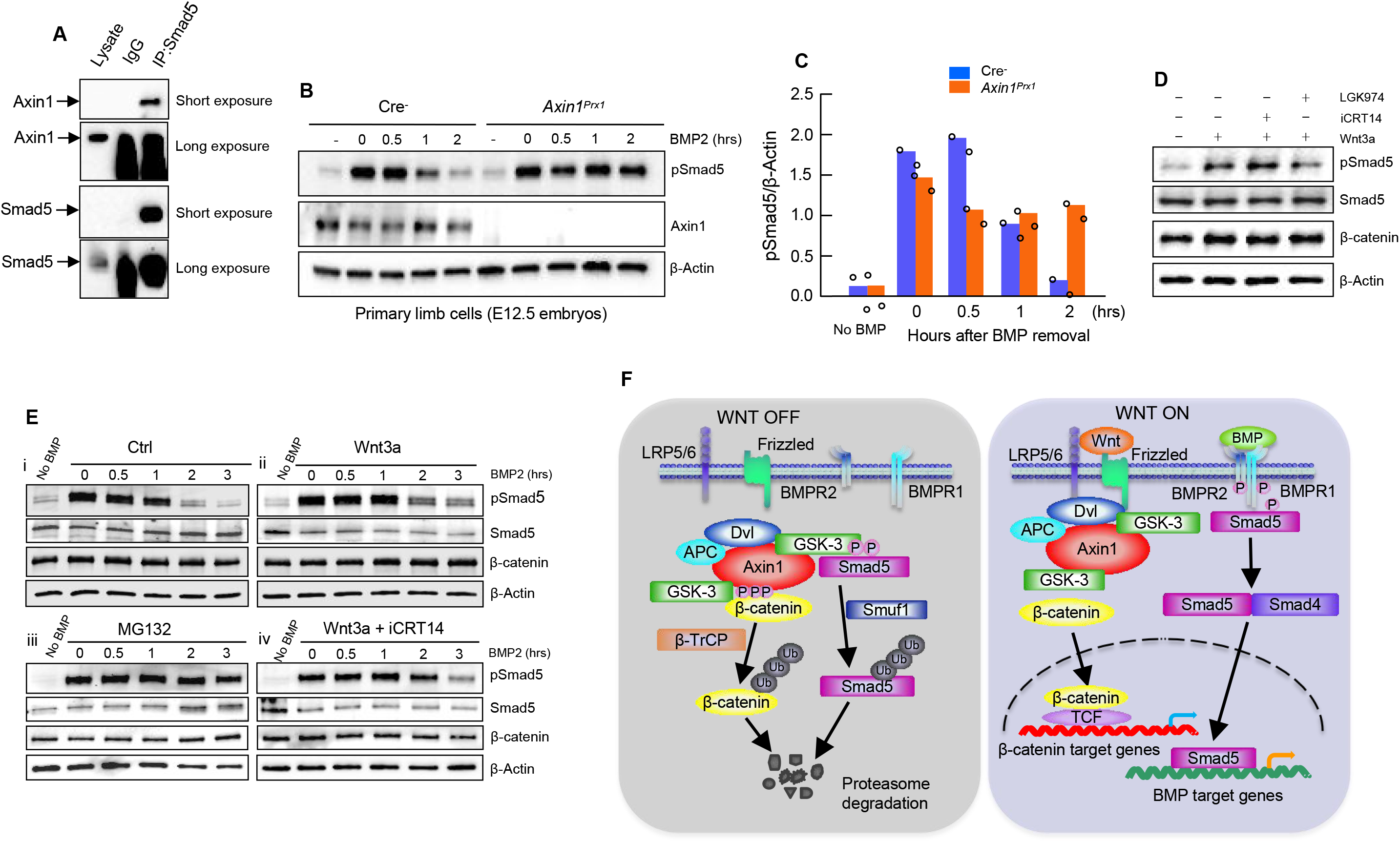
Axin1 regulates BMP signaling through increasing the degradation of pSmad5. (**A**) Interaction of endogenous Axin1 with Smad5 in C3H10T1/2 cells. Co-immunoprecipitation (co-IP) assay was performed using the anti-Smad5 antibody followed by Western blot analysis using the anti-Axin1 antibody. (**B**) Pulse-chase experiments were performed showing that the duration of endogenous pSmad5 were extended in *Axin1*-deficient limb cells. Primary limb cells from Cre^-^ and *Axin1^Prx1^* KO embryos (E12.5) were treated with BMP2 for 0.5 hour, followed by BMP2 wash-out. Cell lysates were extracted at different time points and analyzed by Western blot analysis using indicated antibodies. (**C**) Quantification of the pSmad5 band. (**D**) Wnt3a regulated Smad5 phosphorylation is independent of β-catenin. C3H10T1/2 cells were treated with Wnt3a for 1 hour in the absence or presence of iCRT14 or LGK974 and then harvested for Western blot analysis using indicated antibodies. (**E**) Wnt3a prolonged the duration of pSmad5 in a β-catenin-independent manner. C3H10T1/2 cells were stimulated by BMP2 for 0.5 hour in the absence (i) or presence (ii) of the Wnt3a, proteasome inhibitor MG132 (iii), or the combination of both Wnt3a and iCRT14 (iv). Cells were harvested at the indicated time points after BMP2 wash-out and total cell extracts were analyzed by Western blot analysis. (**F**) Model of integration of Wnt and BMP signaling pathways by Axin1. In the absence of Wnt stimulation, β-catenin is degraded by the destruction complex. Smad5 is also degraded by the Axin1 destruction complex (left panel). In the presence of Wnt ligands or in the absence of Axin1, β-catenin and pSmad5 degradation are inhibited, resulting in activation of both β-catenin and BMP/pSmad5 signaling (right panel).

## Discussion

FH is the most common deficiency of long bone and the pathologic mechanisms of FH is currently unknown. Our present study clearly demonstrated that Axin1/Axin2 play a key role in lower limb development and FH pathogenesis through regulation of both β-catenin and BMP signaling. Defects in Axin1/Axin2 signaling leads to phenotypes resembling FH and PFFD diseases. FH is a birth defect where part of or entire fibulae are missing, as well as associated with limb length discrepancy and foot and knee deformities. FH is a rare genetic disorder, occurring in about 1 in 40,000 live births. In the present studies, we found that the FH phenotype was observed in *Axin1^Prx1^* KO mice, but not in *Axin1^Sox9ER^*, *Axin1^Col2^* and *Axin1^Osx^* KO mice, suggesting that specific deficiency of *Axin1* in *Prx1*-expressing cell population is responsible for the FH development. In addition, we also found that β-catenin or BMP inhibitor can only rescue the FH phenotype during E9.5-E12.5 developmental stages. These findings suggest that FH disease may be caused by specific upregulation of β-catenin or BMP signaling in limb mesenchymal cells during early stage of skeletal development.

The major defect observed in *Axin1^Prx1^* KO mice is the presence of various fibular deficiencies. Some of *Axin1^Prx1^* KO mice show complete or almost complete absence of fibula. The other *Axin1^Prx1^* mice had partial absence of fibulae. All *Axin1/2* dKO mice displayed a complete absence of fibulae. These results indicate that both Axin1 and Axin2 are required for fibular development. As we described above, the partial or total absence of fibulae are the key feature of FH. The *Axin1^Prx1^* KO mice also exhibit tarsal coalition and *Axin1/2* dKO mice show more severe tarsal joint fusion phenotype. The knee joint fusion was also observed in some of *Axin1/2* dKO mice. It is worth noting that the joint fusion phenotype observed in *Axin1^Prx1^* KO mice and *Axin1/2* dKO mice are similar to those found in *noggin* mutant mice (Brunet et al., 1998). Noggin is a secreted BMP antagonist. In addition, the *Axin1/2* dKO mice had femoral defects resembling to PFFD disease (Gillespie et al., 1983).

As Axin1 is a key negative regulator of canonical Wnt pathway and β-catenin is a major downstream mediator of Wnt signaling, we determined genetic interactions between Axin1 and β-catenin during lower limb development. Indeed, the defects in *Axin1^Prx1^* KO mice are significantly rescued by the deletion of one allele of the *β-catenin* gene, *Ctnnb1*. Furthermore, we found that β-catenin inhibitor iCRT14 significantly reversed the FH phenotype in *Axin1^Prx1^* KO mice. These results strongly suggest that Axin1 controls lower limb development through canonical β-catenin pathway.

As mentioned above, we noted that joint fusion phenotype in *Axin1^Prx1^* KO mice could be found in BMP-antagonist mutant mice (Brunet et al., 1998). These findings indicate that BMP signaling may be a downstream signaling pathway of Wnt/Axin1 signaling during limb development. In this study we found upregulation of BMP signaling in *Axin1^Prx1^* KO mice. More importantly, we found that inhibition of BMP signaling efficiently rescued the FH phenotype observed in *Axin1^Prx1^* KO mice. These results indicate that BMP signaling pathway is another key downstream effector of Wnt/Axin1 signaling during limb development. Therefore, it is clear that Axin1 controls lower limb development through both canonical β-catenin and BMP signaling pathways.

It is known that the Wnt and BMP signaling pathways coordinately govern many developmental processes. However, the mechanism of how the two pathways is integrated with each other during skeletal development remains elusive. We found that pSmad5 stability is significantly prolonged in *Axin1*-deficient cells. And we demonstrated that increased pSmad5 stability in *Axin1*-deficient cells is independent from β-catenin activity. Next, we found Wnt indeed regulates Smad5 phosphorylation in C3H10T1/2 cells, in consistent with previous reports (Fuentealba et al., 2007). Importantly, we demonstrated that this regulation is independent from β-catenin signaling. In addition, we confirmed that Axin1 directly interacts with Smad5 in C3H10T1/2 cells. Taken together, we conclude that Axin1 is not only a key negative regulator of β-catenin, but also simultaneously a critical negative regulator of Smad5.

Here, we propose a mechanism by which integration between Wnt and BMP signaling pathways by Axin1 controls limb development and homeostasis. In the absence of Wnt stimulation, β-catenin and Smad5 are degraded by the same destruction complex through interaction with Axin1. In the presence of Wnt ligands or loss of Axin1, both β-catenin and Smad5 degradation is inhibited, resulting in activation of β-catenin and Smad5 signaling (Figure 5F). It is well established that LRP6 recruits the active destruction complex to form the signaling complex with Wnt stimulation, which results in inactivation of the destruction complex, leading to β-catenin stabilization (Li et al., 2012; Kim et al., 2013). However, how Wnt ligands increase the stability of pSmad5 though Axin1 complex remains unknown. It has been reported that Wnt inhibits GSK-3β activity and prolongs the duration of BMP/pSmad1, leading to increased stability of Smad1 (Fuentealba et al., 2007).

It is worth mentioning that BMP signaling is downstream of β-catenin signaling during lower limb development, although we did find that Axin1 could directly interact with Smad5 because β-catenin inhibition could efficiently rescue FH phenotype in *Axin1^Prx1^* KO mice. If Axin1 could regulate lower limb development through interacting with BMP signaling molecules, completely independent from β-catenin signaling, inhibition of β-catenin signaling should not rescue the FH phenotype observed in *Axin1^Prx1^* KO mice.

In summary, in the present studies, we demonstrated that Axin1 and Axin2 are key regulators of lower limb development. Therefore, we hypothesize that transient inhibition of Axin1 or Axin1/2 signaling or transient activation of β-catenin or BMP signaling during early skeletal development may be the cause of FH and PFFD diseases. Our study may have significant impacts on the diagnosis and treatment of these diseases.

## Materials and methods

### Mice

Conditional Axin1 loss-of-function mutant mice (*Axin1^Prx1^*) were generated by intercrossing double heterozygous for a floxed *Axin1* allele and the *Prx1-Cre* transgenic allele (*Axin^flox/wt^;Prx1-Cre^+/-^*) (Logan et al., 2003) with homozygous floxed *Axin1* (*Axin^flox/flox^*) mice. Generation and characterization of *Axin1^flox/flox^* mice was previously reported (Brault et al., 2011). Both males and females of *Axin1^flox/flox^* mice are viable and fertile, and did not present any recognizable phenotype. Axin2 KO mice were obtained from the laboratory of Dr. Wei Hsu (University of Rochester, NY) and have been described previously (Yu et al., 2005; Yan et al., 2009). Mice with floxed *Ctnnb1* (*β-catenin^flox/flox^*), in which exons two to six of the *Ctnnb1* gene are located within *loxP* sites (Brault et al., 2001), were obtained from Jackson Laboratory. All the mice were maintained under standard laboratory conditions with a 12-h light/dark cycle. All animal procedures were approved by the Institutional Animal Care and Use Committee of Rush University Medical Center and all experimental methods and procedures were carried out in accordance with the approved guidelines.

For treatment with Wnt/β-catenin or BMP inhibitors, *Axin^flox/flox^* mice were crossed with *Axin1^Prx1^* mice. The cross rendered 50% Cre-positive mice (*Axin1^Prx1^*) and 50% Cre-negtive control mice (*Axin^flox/flox^*). The pregnant mice were injected with single dose of β-catenin inhibitor iCRT14 (2.5 mg/kg body weight, i.p. injection) or LGK974 (1.0 mg/kg body weight, i.p. injection) or BMP inhibitor dorsomorphin (2.5 mg/kg body weight, i.p. injection) at E9.5, respectively. The mice were sacrificed at E18.5 or 6-week-old. Dorsomorphin and iCRT14 were purchased from Tocris and LGK974 was purchased from Selleck.

### Radiographic and microcomputed tomography (μCT) analyses

Radiographs of mouse skeleton were taken after sacrifice of the animal using a Faxitron Cabinet X-ray system (Faxitron X-ray, Wheeling, IL). For micro-CT analysis, bones were fixed in 10% buffered formalin, stored in 70% ethanol and scanned using a Scanco VivaCT 40 system conebeam scanner (Scanco Medical, Bassersdorf, Switzerland).

### Skeleton preparation and histology

Skeleton preparation and Alizarin red/Alcian blue staining were performed as described (McLeod, 1980). Briefly, mice were sacrificed using carbon dioxide, skinned, eviscerated, and fixed in 95% ethanol. Samples were placed in acetone to remove residual fat. Then the skeletons were stained by Alizarin red/Alcian blue. The stained skeletons were sequentially cleared in 1% potassium hydroxide. The cleared skeletons were transferred into 100% glycerol. For histology, samples were fixed in 10% formalin, decalcified, and embedded in paraffin. 3 m sections were collected and stained with Alcian blue/hematoxylin and eosin (H&E) and Safranin O/fast green following standard procedure.

### Cell culture

For the primary limb cells, forelimb and hindlimb buds of 12.5 dpc *Prx1-Cre* negative or Cre positive homozygous *Axin1* flox embryos were collected in Hanks Balanced Salt Solution (HBSS, Sigma) and digested with 0.1% trysin, 0.1% collagenase in HBSS at 37°C for 15 min. The cells were disassociated through vigorous pipetting, spun down, resuspended in DMEM-F2, 10% FCS and plated in 6-well plates at 2×10^7^ cells per well. Medium was changed daily. C3H10T1/2 mesenchymal progenitor cells were cultured in DMEM supplemented with 10% fetal bovine serum (FBS) and 1% v/v penicillin/streptomycin. Cells were grown at 37°C and 5% CO_2_ in a humidified incubator.

Prior to treatment with BMP2 (25 ng/ml, R&D systems), Wnt3a (100 ng/ml, R&D systems, cells were incubated in serum-free medium for 20 hours. Chemical inhibitors iCRT14 (25 M, Tocris) and MG132 (10 M, Sigma) were added 1 hour prior to BMP2 pulse. LGK974 (20 M, Selleck) was added 6 hours prior to BMP2 or Wnt3a addition.

### Western blot analysis

Cells were lysed with RIPA lysis buffer and protease inhibitor cocktail (sigma #P8340). Protein concentration was determined by Pierce protein assay reagent (Thermo Fisher Scientific #1861426). Protein lysates were boiled in sample buffer (Bio-Rad #1610737). Protein samples were resolved on 10% precast gels (Bio-Rad #4568036) and transferred onto Nitrocellulose membranes (Bio-Rad #1704159). Membranes were blocked in 5% blocking buffer and followed by incubation with primary antibodies and then detected with secondary antibody. The primary antibodies were specific for Smad5 (sc-7443), pSmad1/5 (CST #13820), β-catenin (sc-7963), Axin1 (Millipore #05-1579), Msx2 (sc-15396), Axin2 (ab109307). Secondary antibodies were either HRP-conjugated goat anti-rabbit IgG (Bio Rad #1706515) or rabbit anti-mouse IgG (Bio Rad #1706516) and were revealed with Clarity^™^ western ECL substrate (Bio Rad #1705061). Blots were exposed and scanned by ChemiDoc^™^ xRS+ system (Bio Rad).

### Co-immunoprecipitation

The C3H10T1/2 mesenchymal progenitor cells were washed and collected with cold PBS, lysed in cold lysis buffer containing 50 mM Tris (pH 7.4), 150 mM NaCl, 1 mM EDTA, 1% Nonidet P-40, 10% Glycerol, 0.5 mM DTT, protease inhibitor cocktail tablets (EDTA-free) (Roche) and phosphatase inhibitor cocktail tablet (Roche). The cell lysates were precleared with IgG-agarose beads (Sigma) for 8 hours at 4°C.

Immunoprecipitation (IP) was carried out by incubating the cell lysates with anti-Smad5 antibody (Cell Signaling), robbit IgG immobilized on Protein G Plus-Agarose bead (Santa Creuz) at 4°C overnight. The immunocomplexes were pelleted and washed with cold lysis buffer six times. The proteins were released from beads by boiling in SDS sample buffer, and the samples were analyzed by western blotting.

### Real-time PCR

Total RNA was isolated from cells or tissues by using RNeasy Mini Kit (QIAGEN). Reverse transcription was performed with iScript^™^ Reverse Transcription Supermix Kit (Bio-Rad). Realtime PCR was performed by a Bio-Rad SYBR Green kit and iCycler. Primers are listed in Table 1.

**Table 1.**
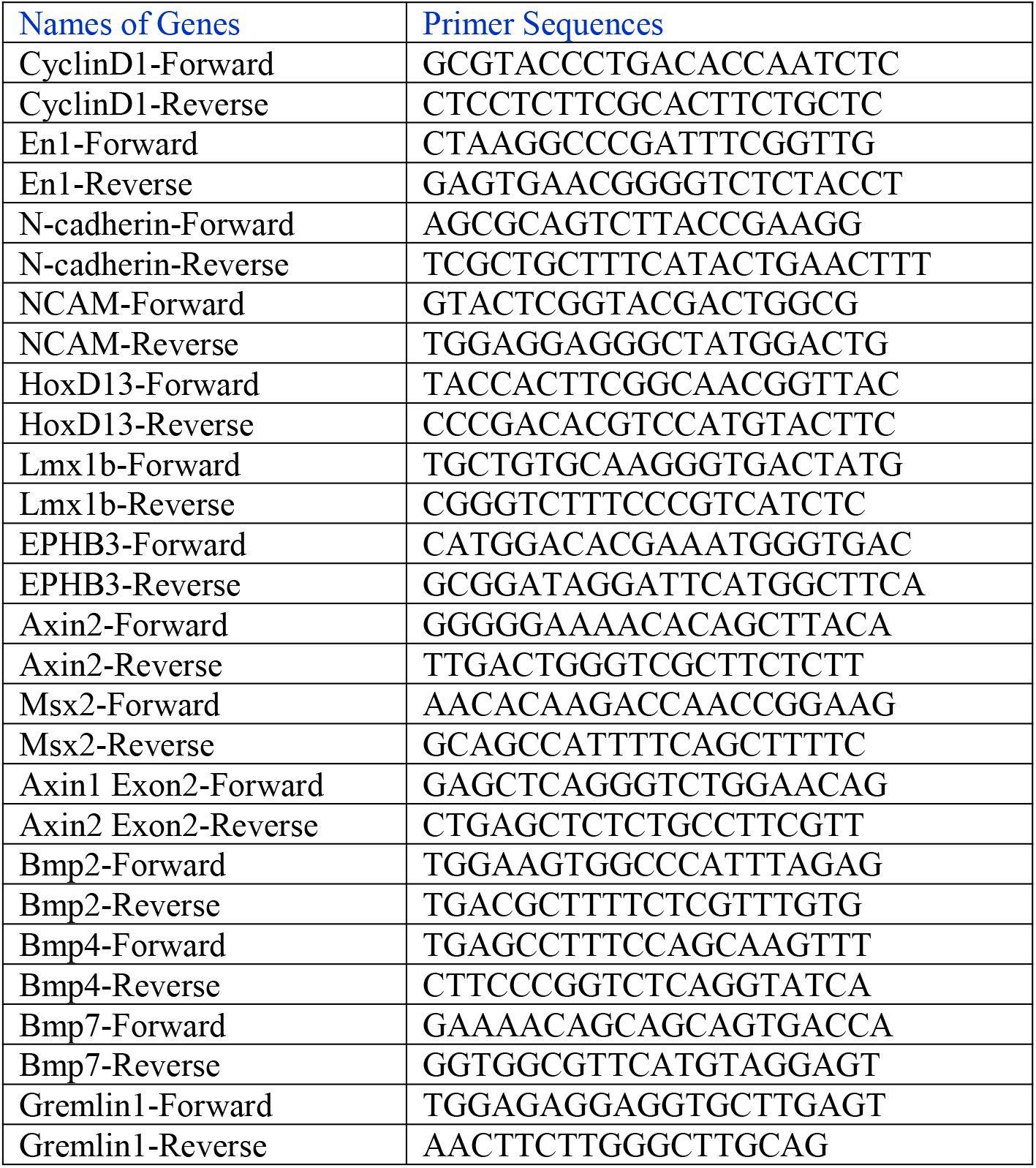
Names of genes and the primer sequences

## Acknowledgments

We would like to express our gratitude to Ms. Lily Yu for her help on processing and staining histological samples.

## Author contributions

D.C. conceived the project and supervised the research. D.C. and R.X. designed the experiments and wrote the manuscript with input from all authors. R.X., D.Y., and Zeng Z. collected and analyzed the data for the project. Q.K., Q.J., Zhenlin Z., G.X., and L.C. provided input for data analysis and manuscript revision. All authors read and commented on the manuscript.

## Data and materials availability

All data needed to evaluate the conclusions in the paper are present in the paper. Additional data related to this paper may be requested from the authors.

